# Inflexible Updating of the Self-Other Divide During a Social Context in Autism; Psychophysical, Electrophysiological, and Neural Network Modeling Evidence

**DOI:** 10.1101/2020.11.02.364836

**Authors:** Jean-Paul Noel, Renato Paredes, Emily Terrebonne, Jacob I. Feldman, Tiffany Woynaroski, Carissa J. Cascio, Peggy Seriès, Mark T. Wallace

## Abstract

Autism spectrum disorder (ASD) is a heterogenous disorder predominantly characterized by social and communicative differences, but increasingly recognized to also alter (multi)sensory function. To face the heterogeneity and ubiquity of ASD, researchers have proposed models of statistical inference operating at the level of ‘computations’. Here, we attempt to bridge both across domains – from social to sensory – and levels of description – from behavioral computations to neural ensemble activity to a biologically-plausible artificial neural network – in furthering our understanding of autism. We do so by mapping visuo-tactile peri-personal space (PPS), and examining its electroencephalography (EEG) correlates, in individuals with ASD and neurotypical individuals during both a social and non-social context given that (i) the sensory coding of PPS is well understood, (ii) this space is thought to distinguish between self and other, and (iii) PPS is known to remap during social interactions. In contrast to their neurotypical counterparts, psychophysical and EEG evidence suggested that PPS does not remap in ASD during a social context. To account for this observation, we then employed a neural network model of PPS and demonstrate that PPS remapping may be driven by changes in neural gain operating at the level of multisensory neurons. Critically, under the anomalous excitation-inhibition (E/I) regime of ASD, this gain modulation does not result in PPS resizing. Overall, our findings are in line with recent statistical inference accounts suggesting diminished flexibility in ASD, and further these accounts by demonstrating within an example relevant for social cognition that such inflexibility may be due to E/I imbalances.

## Introduction

Autism spectrum disorder (ASD) is a neurodevelopmental condition characterized by altered social interactions, repetitive behaviors and restricted interests, and differences in language and communication skills. The current diagnosis rate is approximately 1 in 59 children in the United States (Xu et al., 2018), and in addition to growing in pervasiveness, it is a condition growing in scope. Beyond the established core features in the social domain, sensory processing differences are increasingly recognized in ASD (Robertson & Baron-Cohen, 2018). In fact, atypical sensory responses are now part of the ASD diagnostic criteria (DSM-V).

Two overarching themes have emerged in the study of sensory function in ASD. The first of these comes from animal model work, and highlights a marked hyper-sensitivity of neurons during weak sensory stimulation in mouse models of autism (e.g., Goncalves et al., 2013; Orefice et al., 2016). In principle, these changes could be a property of the neurons themselves, but they are more widely considered to be a circuit property (Chen et al., 2020) reflecting changes in the balance of excitation and inhibition (E/I imbalance, Rubenstein & Merzenich, 2003; Lee et al., 2017). The second theme, derived largely from the human literature, suggests that while individuals with ASD may outperform their neurotypical counterparts in local sensory processing (e.g., Joseph et al., 2009), they present with decreased holistic processing (i.e., weak central coherence; Happe & Frith, 2006). A similar conclusion applies to the ability to integrate information across different sensory modalities, with a growing body of evidence pointing to atypical multisensory integration in ASD (see Baum et al., 2015, and Wallace et al., 2020, for reviews, but also Zaidel et al., 2015, for opposing evidence).

Now, bridging across the core features deficits that characterize ASD, their phenotypic-level sensory processing (dis)abilities (e.g., multisensory integration), and their putative underlying neural implementation level causes (e.g., E/I imbalance), is a notorious challenge. Arguably, this endeavor requires both (1) the appropriate computational tools to bridge across levels of description (i.e., from neurons to behavior) and, (2) an experimental paradigm that is germane to both social and sensory processing.

Regarding the former, the recent rise of computational psychiatry reflects an attempt to both broadly ascribe the heterogeneity of psychiatric disorders (including ASD) to underlying algorithmic anomalies, as well as to bridge across levels of description (e.g., Huys et al., 2016). In autism, much of this current effort is centered around weaknesses in the ability of those with ASD to perform and update statistical inferences (Pellicano & Burr, 2012; Palmer et al., 2017; Karvelis et al., 2018) – a computation which is ubiquitous across brain networks and fundamental for perception (e.g., Pitkow & Angelaki, 2017). However, these studies are limited in their ability to link across levels of description, e.g., from phenomenology to neural circuitry. Regarding the latter, much of the work bridging between perceptual and social deficits in ASD has focused on establishing correlations between multisensory (i.e., audio-visual) temporal perception and social-communicative abilities (e.g., language comprehension; Woynaroski et al., 2013; Stevenson et al., 2014; see Wallace et al., 2020). The linkage from language comprehension to neurons, however, may be a bridge too far for current computational tools.

In the current study, we take advantage of a spatial multisensory paradigm that offers the opportunity to bridge from sensation to social cognition. Namely, it is well established that there is a network of multisensory (e.g., visuo-tactile) neurons located in posterior parietal and ventral pre-motor cortices that map the space immediately surrounding and adjacent to the body – the peri-personal space (PPS; Rizzolatti et al., 1981, 1983, Shelton et al., 1990; Ladavas et al., 1998; Serino, 2019). This space is thought to delineate the space of the self vs. that of other agents (Blanke, 2012, Noel et al., 2015; Salomon et al., 2017), and both physiological recordings in monkeys (Ishida et al., 2010) and psychophysical tasks in healthy humans (Teneggi et al., 2013; Pellencin et al., 2017) have suggested that the PPS remaps as a function of social context (e.g., quality of social interaction or even perceived morality of others). PPS has been reported to be smaller in individuals with ASD than in their neurotypical counterparts (Mul et al., 2019; Noel et al., 2020), yet its reshaping based on social context has not been studied. Further, biologically-plausible neural network models of PPS have been proposed (Magosso et al., 2010a, 2010b; Serino et al., 2015a), and these models are able to recapitulate the basic properties of PPS encoding (e.g., visual or auditory facilitation of tactile processing when the former is near but not far from the body). However, these models have yet been brought to bear on questions of clinical interest in ASD, or to the issue of the malleability of PPS based on social context.

We map PPS in ASD both via psychophysics and electroencephalography (EEG). The neural recordings provides direct evidence for true multisensory integration (visuotactile) in individuals with ASD that is modulated by the distance between constituent cues (visual and tactile). These approaches suggested an inflexible mapping of PPS and an immutable degree of multisensory integration in ASD regardless of social context. Neural network modeling suggests that this inflexibility may be due to a gain modulation provided by the social context being less impactful within the E/I regime characterizing ASD.

## Materials and Methods

### Participants

A total of 50 participants (20 females, 30 males, mean age = 20.8 ± 3.34 years) took part in this experiment. Of these, 20 (8 females, mean age = 21.3 ± 4.5 years) were individuals previously diagnosed by a clinician practitioner as on the autism spectrum according to the diagnostic criteria of the DSM-5, and further evaluated and confirmed as individuals with ASD by a research-reliable clinician using the Autism Diagnosis Observation Schedule (ADOS-2,; Lord et al., 2012). The other 30 participants were neurotypical control individuals group-matched for age, gender, and IQ to the ASD group (all p > 0.18). Individuals within the control group did not have a diagnosis of ASD or any other psychiatric or neurologic disorder. All participants completed the Social Responsiveness Scale (SRS; Constantino et al., 2003) and as expected ASD participants scored higher (mean ± sem, 95.15 ± 6.30) than the neurotypical participants (44.40 ± 4.29, p = 1.59 x 10^−8^). Participants gave consent in according to the Declaration of Helsinki before taking part in the experiment, and all protocols were approved at the Institutional Review Board at Vanderbilt University.

### Materials and Apparatus

Visual and tactile stimuli were driven via a micro-controller (SparkFun Electronics, Redboard, Boulder CO) and direct serial port communication under the control of purpose written MATLAB (MathWorks, Natick MA) and Arduino (Arduino™) scripts. Visual stimuli were a flash of light from a red LED (3 mm diameter, 640 nm wavelength), while tactile stimuli consisted of vibrotactile stimulation administered via a mini motor disc (10mm diameter, 2.7mm thick, 0.9 gram, 5V, 11000 RPM). These stimuli were 50 ms in duration (square-wave, onset and offset <1 ms, as measured via oscilloscope). The LEDs and vibrotactile motor were mounted in an opaque enclosure where 30 LEDs sequentially protruded above the enclosure every 3.3 cm (in depth) and counted with a hand rest immediately adjacent to the first LED (see Noel et al., 2018a for a similar apparatus). In the current study LEDs number 5, 8, 11, 14, and 17 were utilized, corresponding to visuotactile depths of 13.2cm, 23.1cm, 33.0cm, 42.9cm, and 52.8cm. Visuotactile stimuli consisted of the synchronous presentation of the visual and tactile stimuli described above.

### Procedure

Participants were seated in a light- and sound-controlled room in which they performed a tactile reaction time (RT) task to a tactile stimulation administered on their left index finger. Responses were given by button-press with their right index finger. Trials could be visuotactile (i.e., experimental trials; VT), tactile (i.e., baseline; T), or visual (i.e., catch; V) trials. Visual trials were ‘catch trials’ in that they did not require a motor response, while tactile trials were ‘baseline trials’ as these permitted us to gauge tactile RTs in the absence of visual inputs, and hence determine whether a multisensory effect was present as a function of visuo-tactile distance. Visuotactile and visual trials were presented at 5 different distances (D1 through D5 = 13.2cm, 23.1cm, 33.0cm, 42.9cm, and 52.8cm, **Figure 1A**). Within each block 132 trials were presented; 20 VT trials at each of the 5 distances, 4 V trials at each of the 5 distances, and 12 T only trials. Trial type was randomized within blocks, and inter-trial interval was random between 1500-2250ms (uniform distribution). Participants completed 8 or 10 blocks (total of 1056 to 1320 trials) according to time constraints. Block duration was ~6 minutes. Half of the blocks were ‘non-social’, and half were ‘social’. During ‘social’ blocks an experimenter sat facing the participant at a distance of 150 cm with a neutral expression (see Teneggi et al., 2013 for a similar manipulation; **Figure 1A**). The order of blocks was counter-balanced across participants.

**Figure 1.**
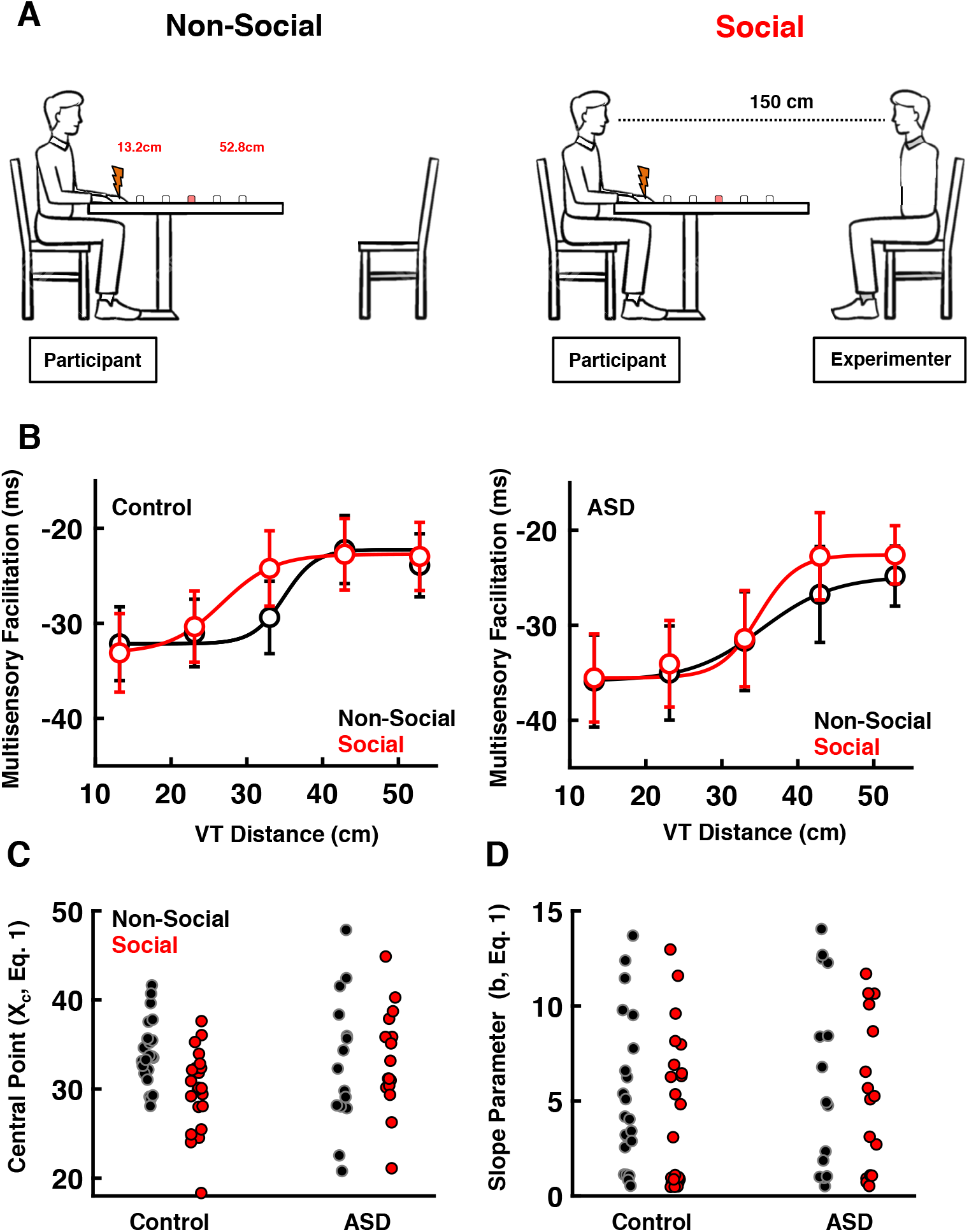
Methods and Behavioral Results. **A) Experimental Setup**. Participants responded as fast as possible to tactile stimuli (orange), which could be paired with visual stimuli (red, example show at third distance) at different distances (D1-D5, 13.2 – 52.8cm). In different blocks, an experimenter would sit facing the participants with a neutral expression (social blocks). In the non-social blocks there was nobody else in the experimental room. **B) Visuo-tactile reaction times as a function of visuo-tactile distance**. Both for neurotypical control (left) and ASD (right) participants, reaction times were further facilitated when visual stimuli were near the body, demonstrating a PPS effect. The facilitation was well expressed in the majority of participants by a sigmoidal function in both the non-social (black) and social (red) blocks. The fit shown is to the average RTs across participants, and not the average fit. **C) Extracted Central Point**. PPS become smaller in neurotypical control (left) but not ASD (right) participants during the social blocks. **D) Extracted Slope Parameter**. The gradient separating the space where visual presentation facilitated vs. not tactile reaction times did not change in either group as a function of social context, and instead was characterized by a marked inter-participant variability.

### Behavioral Analyses

Tactile RTs were measured as the time elapsed from touch onset to button response. These responses were coalesced as a function of sensory modality (unimodal tactile vs. multisensory visuo-tactile), distance (D1-D5), and social context (social vs. non-social). Then, the median RT was computed for each participant and condition. As a preliminary analysis, we performed a paired t-test across sensory modalities in order to confirm that multisensory visuo-tactile trials (regardless of distance) were faster than unisensory tactile trials, and thus as expected visual presentations facilitated tactile responses (Noel et al., 2019). Next, we performed a one-way ANOVA to confirm that tactile facilitation was dependent on the proximity of visual presentation; namely, to query whether as a whole participants demonstrated a PPS effect (e.g., Pfeiffer et al., 2018). Lastly, on a participant-per-participant basis we fit RTs to a sigmoidal function (**Eq. 1**),

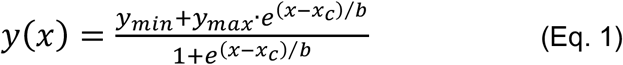

from which we extracted the central point of the sigmoidal (*x_c_*, in Eq. 1, representing the boundary of PPS) and a parameter inversely proportional to its slope at the central point (*b* in Eq. 1, characterizing the gradient of PPS representation; see Noel et al., 2018, 2020, for a similar approach). The central point and slope of the sigmoidal fit were contrasted via a mixed ANOVA as a function of social context (i.e., social vs. non-social), the participant group (i.e., control vs. ASD), and the interaction of these variables to describe the shape (size and gradient) of PPS encoding. An a priori cutoff r-square >0.25 was applied, excluding data at the participant level if this threshold was not reached on both the social and non-social conditions. This resulted in 9/50 (18%) participants being excluded. To ascertain whether results held on the sample as a whole, we additionally employed the Spearman-Karber method (see Bausenhart et al., 2018 for a tutorial review, and see Masson et al., 2020 for prior application of this method to PPS data) to estimate the central point and slope of the psychometric functions without requiring a fitting procedure.

### EEG Recording and Preprocessing

Continuous EEG was recorded from 128 electrodes with a sampling rate of 1000Hz (Net Amps 400 amplifier, Hydrocel GSN 128 EEG cap, EGI systems Inc.) and referenced to the vertex (Cz). Electrode impedances were maintained below 40 kΩ throughout the recording procedure. Data were acquired with NetStation 5.1.2 and further pre-processed using MATLAB and EEGLAB (Delorme and Makeig, 2004). Continuous EEG data were notch filtered at 60 Hz and bandpass filtered from 0.1 Hz to 40 Hz using an 8th order bi-directional zero-phase infinite impulse response (IIR) filter. Epochs from 500 ms before to 1000 ms after stimuli onset were extracted and divided according to experimental condition. Artifact contaminated trials (e.g., eye-blinks, 11.9% ± 4.5%) and bad channels (1.2% ± 0.7%) were identified and removed through a combination of automated rejection of trials in which any channel exceeded ± 100 mV and rigorous visual inspection. Data were then recalculated to the average reference and bad channels were reconstructed using spherical spline interpolation (Perrin et al., 1987). Lastly, data were baseline corrected for the pre-stimuli period (−200 to 0 ms post-stimuli onset).

### EEG Analyses

We quantified the global electric field strength present throughout the recording montage using global field power (GFP; Lehman & Skrandies, 1980). This measure corresponds to the standard deviation of the trial-averaged voltage values across the entire electrode montage at a given time point, and represents a reference- and topographic-independent measure of evoked potential magnitude (Murray et al., 2008). Further, GFP is additionally a data-reduction technique given that it summarizes 128 distinct time-series (i.e., electrodes) into a singular one, thus reducing the multiple comparisons problem in EEG and the possibility for false positives.

In a first pass, we calculated the GFPs for each participant, as well as for the entire sample of participants (i.e., grand average) for tactile, visual, and visuotactile conditions separately while collapsing across distances (see **Figure 2**). Time-resolved t-tests against zero were performed at each time-point from 200 ms pre-stimuli presentation to 1000 ms post-stimuli onset in order to ascertain whether reliable evoked potentials were generated (to V, T, and VT stimuli). To account for the inherent auto-correlation problem in EEG, we set alpha at < 0.01 for at least 10 consecutive time points (Guthrie & Buchwald, 1991; see Simon et al., 2017, and Cascio et al., 2015, for a similar approach). Next, to ascertain whether a veritable multisensory effect existed (i.e., nonlinearity between the co-presentation of V and T information vis-à-vis their presentation in isolation; e.g., Stein & Stanford, 2008) we created artificial visuo-tactile summed responses (hereafter, “summed” or “sum”) by adding the participant-level average responses to V and T. GFP was then calculated for this summed response and contrasted to the GFP of the multisensory visuotactile condition (or “paired” response; see Cappe et al., 2010 for a similar approach). Indeed, as GFP is by definition positive, an advantage of utilizing this method within a multisensory context is that super- and sub-additivity indices may be measured with no ambiguity due to polarities (e.g., Sperdin et al., 2010). Having established that the co-presentation of visual and tactile information resulted in a multisensory effect, and having identified the time-period and spatial cluster driving this effect, we next computed the multisensory effect (i.e., difference in voltage between summed and paired responses within the spatio-temporal window of interest) for each visuo-tactile distance, social context, and participant group. These latter variables were contrasted via a 5 (distance) by 2 (social context) by 2 (participant group) mixed ANOVA.

**Figure 2.**
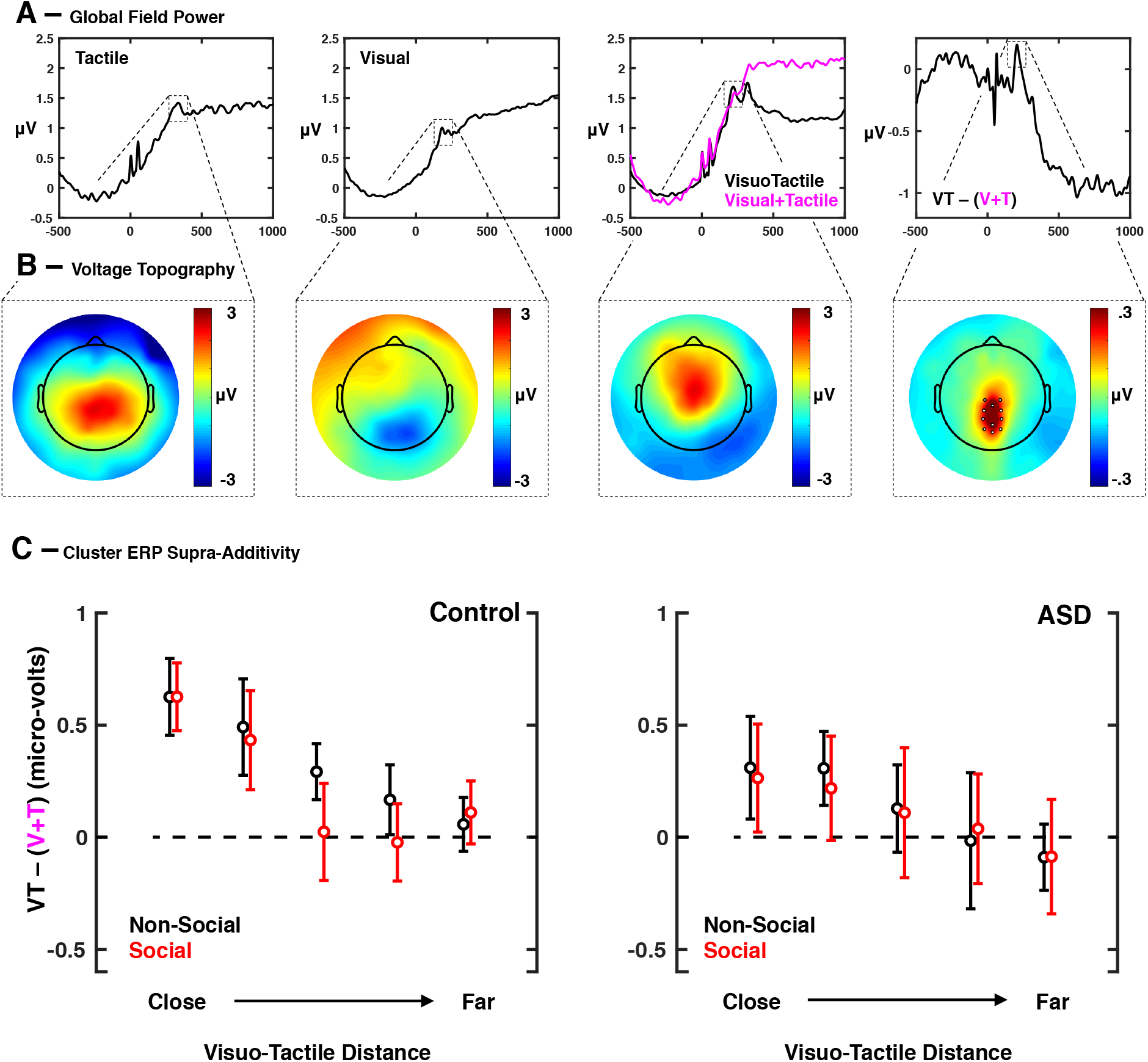
Electroencephalography Results. **A) Global Field Power as a function of sensory modality**. Results show clear evoked responses for tactile (leftmost), visual (2^nd^ panel), and visuo-tactile (black, 3^rd^ panel) stimuli presentations. More importantly, the visuo-tactile response shows true multisensory integration, being stronger than the artificially summed visual+tactile response (magenta). Rightmost panel shows the difference wave between the paired and summed response. **B) Topography of responses and difference wave (rightmost)**. The topography of responses (nose at the front) is indicative of tactile (leftmost), visual (2^nd^ panel), and visuotactile (3^rd^ panel). The rightmost panel shows that the supra-additive multisensory effect is driven by electrodes in centro-occipital areas, between unisensory visual and somatosensory areas. **C) Voltages during supra-additive period within the cluster driving the GFP effect**. The difference in voltage between the paired (VT) and summed (V+T) conditions show a clear multisensory integration effect, and one that is dependent on spatial disparity between V and T. Most interestingly, social context (black = non-social, red = social) modified the degree of multisensory integration in controls (left) but not ASD (right) individuals.

### Neural Network Modeling

To examine putative circuit-level mechanisms subserving the behavioral and physiological results observed, we implemented and extended a well-established biologically-plausible neural network model of PPS (Magosso et al., 2010a, b; Serino et al., 2015a; Noel et al., 2018b, 2020a). The model is composed of non-spiking (rate) neurons whose output is a continuous variable representing the neuron’s firing rate. The network includes two recurrently connected unisensory areas (tactile and visual) projecting onto a third area composed of a multisensory visuo-tactile neuron (see **Figure 3**).

**Figure 3.**
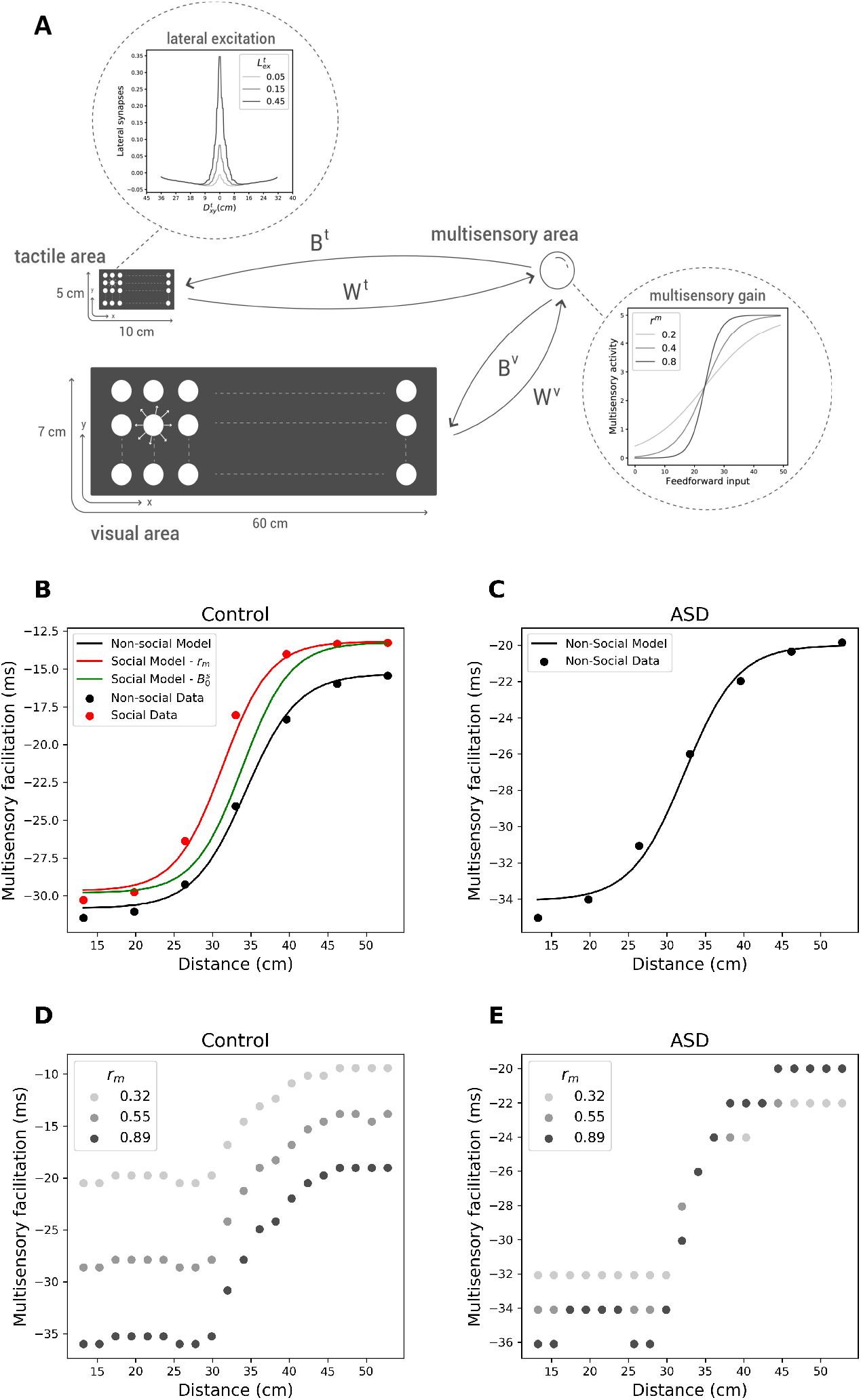
Neural Network Model. **A) Neural Architecture**. The neural network is composed of a tactile area coding for the hand, a visual area coding for near and far space, and a multisensory neuron receiving projections from the unisensory areas, and reciprocally sending feedback projections back to unisensory areas. The output of each neuron is dependent on input-output functions, and the inset to the right shows examples of different gain functions for the multisensory neuron. Within unisensory areas, neurons are laterally connected by a “Mexican-Hat” pattern, with near excitation and far inhibition (see inset to the top). **B) Model Fits to Control Participants**. As a first step we fit multisensory facilitation (y-axis) in reaction time as a function of distance from the body (x-axis) in the neurotypical control and non-social condition (black). Seven distances generated from a sigmoidal with parameters equal to the median experimental parameters were used as observed data. The model is well able to account for observations. Then, we try to explain the impact of the social manipulation by either a change in neural gain at the level of the multisensory neuron (red) or the strength of feedback projects (green). The former approach accounted best for observed data. **C) Model Fits to individuals with ASD**. The strength of excitatory lateral connections were allowed to vary from the non-social control and the non-social ASD model, and this manipulation was well able to account for idiosyncrasies in the shape of PPS in ASD (see main text and simulations in supplementary note). **D) Impact of Multisensory Gain under the Control E/I Regime**. Increasing gain of the multisensory neuron (from light gray to black) increased the size of PPS. **E) Impact of Multisensory Gain under the ASD E/I Regime**. Increasing gain of the multisensory neuron (from light gray to black) did not impact the size of PPS.

Each unisensory area is composed of a matrix of unisensory neurons. The tactile area is composed of 200 neurons (20 x 10 grid) covering a skin portion of 10 cm x 5 cm. The visual area is composed of 1680 neurons (120 x 14 grid), covering a visual space of 60 cm in depth and 7 cm horizontally. These unisensory neurons each have a bivariate Gaussian receptive field through which stimuli are convolved, and cells within each area are reciprocally connected via lateral synapses *L* having a “Mexican-hat pattern” (i.e., near excitation and far inhibition). The specific weights assigned to each synapse are obtained as the difference between two Gaussians (one excitatory and one inhibitory) according to,

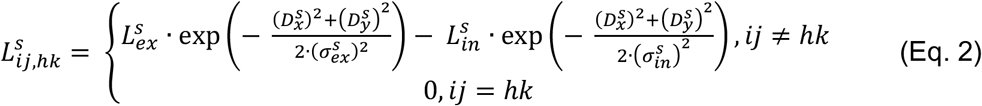

where 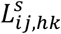 denotes the weight of the synapse from a pre-synaptic neuron at position *hk* to post-synaptic neuron at position *ij*, the superscript *s* can take on *t* (i.e., tactile) or *v* (i.e., visual) as values, and the subscripts *exp* and *in* respectively refer to excitation and inhibition. 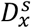 and 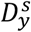 indicate the distances between the pre- and post-synaptic neuron along the horizontal and depth axis of the unisensory area. The null term is included to avoid auto-excitation.

Neurons within both unisensory areas send excitatory feedforward synapses, *W*, to the downstream visuotactile area. In turn, the multisensory neuron sends excitatory feedback connections, *B*, back to the unisensory areas. The feedforward synapses from the tactile neurons to the multisensory one have a uniform value, 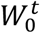. Similarly, the feedback projections from the multisensory neuron to tactile neurons all have strength 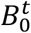. On the other hand, the weights of synapses that connect visual neurons and the multisensory neuron take on their maximal value, 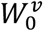 and 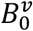, within a certain depth boundary, *Lim*, and then decrease with further depth according to the biexponential functions,

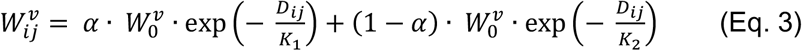

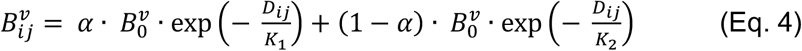

*K*_1_, *K*_2_, and *α* are parameters governing the exponential decay of synaptic weights after the first *Lim* cm.

The overall input, *u*, to a neuron in the network is processed via a first-order temporal dynamics (**Eq. 5**, mimicking the post-synaptic membrane time constant) and a sigmoidal function (**Eq. 6 and 7**, mimicking the neuron’s activation function), generating the neuron’s output activity, *z*(*t*):

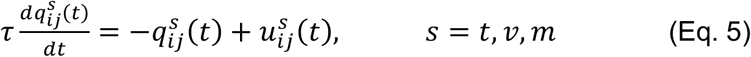

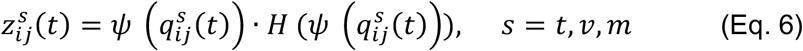

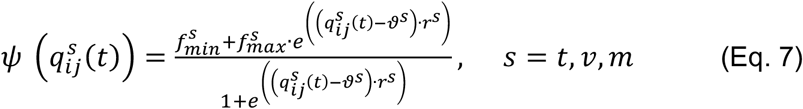

**Eq. 5** holds for both unisensory neurons *s* = *t,v* and for the multisensory neuron 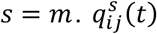 is the state variable of a neuron at a given time step, *t*. *u^s^*(*t*) is the input to the neuron and *τ* the time constant. **Eq. 6** describes a sigmoidal function, *ψ*, being applied to the state variable and this being multiplied by a Heaviside function *H* to avoid negative values in the activity of neurons (similar to a ReLu function in machine learning; Hahnloser et al., 2000). **Eq. 7** describes the sigmoidal activation function; 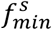 and 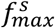 represent the lower and upper saturation of the sigmoidal function, *ϑ^s^* establishes the central value of the sigmoidal function (i.e. the input value at which the output is midway between 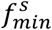 and 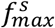) and *r^s^* defines the slope.

The overall input to each unisensory neuron (*u^s^*(*t*), s = t, v) is made up of the external input coming from outside the network (i.e., the stimulus filtered by the neurons’ receptive field, 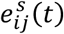, plus the lateral input coming from other neurons in the same area (via weights defined by lateral synapses *L*), and feedback input from the multisensory neuron (via weight defined by the feedback synapses *B*, see **Eq. 8**).

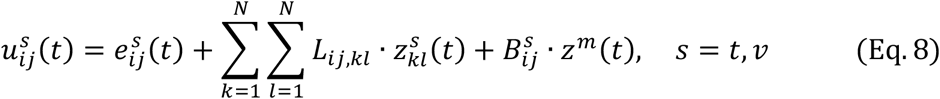

The overall input to the multisensory neuron is made up of the feedforward inputs from the two unisensory areas (via weights defined by the feedforward synapses *W*, see **Eq. 9**).

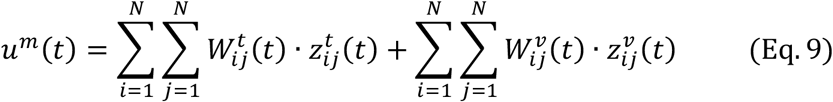

All abovementioned differential equations are solved by numerically employing the Euler integration method with a discrete time-step of 0.4ms. Tactile stimuli within the network were always applied at *x^t^* = 10 *cm* and *y^t^* = 5 *cm*. In contrast, the visual stimuli were always applied at coordinate *y^v^* = 7.5 *cm* and critically *x^v^* changed across trials to take on the true experimental values of visual stimuli during the behavioral experiment.

The reaction time of the network was registered at every distance as the time at which any neuron of the tactile area reached 90% of its maximum state (i.e., 90% of 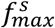). To translate between output of the neural network and human psychophysical reaction times, we also presented the network with tactile-only stimuli, and computed its multisensory facilitation just as we did for the human observers (i.e., difference between unisensory and multisensory reaction times). Then, we matched the time units of the model to real time by linear regression,

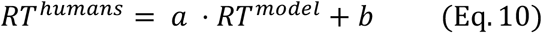

Where *RT^humans^* is the multisensory facilitation measured in humans, *RT^model^* is that measured within the neural network, *a* denotes how many milliseconds correspond to one unit time of the model, and *b* represents the duration of neural processing not captured by the model. Novelty, within the current report we implemented a differential evolution algorithm (see Wormington et al., 1999, Virtanen et al., 2020) to perform model fitting of the PPS neural network. During this fitting *α* was bounded to be positive, and *b* to be negative. To further constrain the neural network model fits, we generated ‘experimental data’ at seven distances (as opposed to the five tested in humans) given the median *y_min_*, *y_max_*, *x_c_*, and *b* (see **Eq. 1**) from the social and non-social conditions in controls and individuals with ASD.

Default parameter values were inherited from previous reports (Magosso et al., 2010a,b) and can be found in **Table S1**. For more detail see **Supplementary Note**.

## Results

### Smaller PPS During Social Context in Neurotypical but not ASD Individuals

Overall participants were very accurate at withholding responses during visual-only catch trials (false positives, 2.3% ± 1.4%) and therefore behavioral analyses focuses on reaction times (RT, see Serino et al., 2015b for a similar approach).

The contrast between tactile and visuo-tactile RTs (regardless of distance) demonstrated a clear multisensory facilitation effect (tactile, 383.3 ms ± 2.9 ms; visuo-tactile, 354.9 ms ± 3.5 ms, t = 10.32, p = 6.85×10^−14^). Moreover, in general the RTs of individuals with ASD (376.6 ms ± 2.8 ms) were slower than that of control individuals (340.5 ms ± 3.1 ms, p = 1.57×10^−4^). Thus, to examine putative space-dependent multisensory effects, we first computed for each participant and visuo-tactile disparity an index of multisensory facilitation (i.e., VT – T); the more negative this value, the stronger the multisensory effect (see Noel et al., 2018, for a similar approach). For both the neurotypical (p = 0.024) and the ASD groups (p = 0.019), facilitation in the detection of a tactile target by a co-presented visual stimulus was space dependent, with the effect being most apparent at smaller visuo-tactile spatial disparities (**Figure 1B**). Thus, both groups showed a modulation of behavior based on the spatial structure of co-presented visual and tactile stimuli, consistent with the encoding of PPS.

To more fully characterize the PPS effect we fit RTs to a sigmoidal function (**Eq. 1**) with its central point and slope at the central point as free parameters. After removal of participants with very poor fits (see *Methods* section), this function closely fit the pattern of RTs (R^2^, 0.78 ± 16.2), and did so equally for the two participant groups (p = 0.43). Although there was a tendency for those with ASD to have a smaller PPS at baseline (ASD, 32.68 cm ± 1.64 ms; neurotypical controls, 34.09 ± 0.67cm, p = 0.18, F-test main effect), in the current dataset this difference failed to reach significance (but see Mul et al., 2019 and Noel et al., 2020, for evidence suggesting smaller PPS in ASD than neurotypical individuals). More importantly, however, to the best of our knowledge the current report is the first to examine the re-sizing of PPS in individuals with ASD as a function of an experimental manipulation – the presence of another individual in this case. In line with Teneggi et al., 2013, the presence of another individual (social condition) appeared to overall shrink our participant’s’ PPS (non-social, 33.39 cm ± 0.89 cm; social, 31.46 cm ± 0.82 cm, p < 0.001, F-test main effect). Critically, this re-sizing was true in control participants (non-social, 34.09 cm ± 1.64 cm; social, 29.66 cm ± 1.28 cm, p = 5.96 ×10^−6^), but not for the ASD group (non-social, 32.68 cm ± 0.67 cm; social, 33.27 cm ± 0.82 cm, p = 0.26; **Figure 1C**). Similar (but nonparametric due to their skewed distribution) analyses on the gradient of PPS (parameter *b* in **Eq. 1**) demonstrated no difference between or within groups (all p > 0.25). The most striking feature of these estimates was their marked variability (see **Figure 1D**).

Lastly, as a confirmatory analysis we estimated again the central point of the psychometric function describing visuo-tactile RTs as a function of visuo-tactile distance via the Spearman-Karber method (see Bausenhart et al., 2018). This method allows for estimating the measures of interest without sigmoidal fitting, and thus we were able to include all participants in this analysis. Results confirmed an interaction between social context and participant group (p = 0.01). This effect was driven by the resizing of PPS within a social setting in the control group (p = 0.021), but not the ASD group (p = 0.12).

### Physiological Marker Suggests Unchanged Multisensory Integration During Social Context in ASD

The behavioral paradigm employed above allows mapping PPS via a task that has been classically used to index multisensory interactions, and suggests that that while PPS in individuals with ASD may be smaller (empirically supported by Mul et al., 2019, Noel et al., 2020, and theoretically suggested by Noel et al., 2017), its starker defining characteristic is its inflexibility. Namely, in contrast to neurotypical individuals, PPS did not remap for ASD individuals during a social context. In this behavioral paradigm, however, participants are instructed not to respond to visual stimuli alone. This experimental design choice was driven by the fact that inclusion of “catch, non-response” trials greatly reduces the expectation bias that results from having to respond on every trial (see Kandula et al., 2017 and Hobeika et al., 2019 for more detail). However, not indexing unisensory visual-only trials impedes from ascertaining whether true multisensory integration occurred, via race (Miller, 1982) or drift-diffusion (Drugowitsch et al., 2014) models, both requiring RTs to every component (e.g., V, T, and VT).

To take a different approach toward assessing true multisensory integration (vs. interactions), we turned to electrophysiology. We sought to assess visuo-tactile multisensory integration as a function of spatial disparity, social context, and clinical diagnosis. In a first step, we computed Global Field Power (GFP, see *Methods*), an index of overall neural response strength. This analysis demonstrated a reliable evoked tactile response beginning at 151 ms post-stimulus onset (p<0.01). This tactile GFP signal peaked at 334 ms post-stimulus onset (**Figure 2A**, leftmost). Visual evoked responses were reliably indexed somewhat later, beginning at 170 ms poststimulus onset (p<0.01), and peaking shortly thereafter at 182 ms post-stimulus onset (**Figure 2A**, 2^nd^ column). Response to combined visuotactile stimulation began 152 ms post-stimulus onset (p<0.01), and peaked at 218 ms post-stimulus onset (**Figure 2A**, 3^rd^ column). In regards to their spatial distribution on the scalp, the tactile response was characterized by a central positivity (**Figure 2B**, leftmost), the visual response was characterized by an occipital negativity (**Figure 2B**, 2^nd^ column), and the visuotactile response was a combination of these topographies, showing both a negativity in posterior electrodes and a positivity pole over fronto-central electrodes (**Figure 2B**, 3^rd^ column).

To determine whether the presentation of visuotactile stimuli elicited a response indicative of multisensory integration, we created an artificial summed (i.e., visual + tactile) response. An actual multisensory (“paired”) response that deviates from this additive prediction is considered a hallmark of multisensory integration (see Stein & Stanford, 2008, for a review, and Bernasconi et al., 2018, and Noel et al., 2019a for a similar approach within the study of PPS). The contrast between the actual and summed GFP demonstrated superadditivity of the multisensory response (V+T<VT) between 153 and 175 ms post-stimulus onset (p < 0.01). This finding was true for both the control (between 154 ms and 170 ms post-stimuli onset, p < 0.01) and ASD (between 158 ms and 174 ms post-stimuli onset, p < 0.01) individuals. While the evoked multisensory response at its peak amplitude was driven by a widespread negativity in posterior electrodes and positivity more anteriorly, the topography of the difference wave was focused with a positivity in centro-posterior electrodes (**Figure 2B**, rightmost, electrodes highlighted in white demonstrated a voltage-difference between the paired and summed response).

Having restricted our time-period of interest to between 158 and 170 ms post-stimulus onset (the union of the time-periods demonstrating a response indicative of multisensory integration in all groups), as well as our montage state space (i.e., spatial window of interest) to those centro-posterior electrodes driving the multisensory effect, **Figure 2B**, rightmost), we next investigated whether this response differed based on visuo-tactile disparity, and how it varied as a function of social context and participant group. We extracted and averaged voltages within the spatio-temporal window defined above for each participant, modality type (T, V, or VT) and distance (in the case of V and VT). We then computed the degree of multisensory superadditivity (VT – (V + T), positive values indicating greater multisensory integration) and performed a 5 (distance) by 2 (social context) by 2 (participant group) mixed ANOVA.

Results demonstrated that the multisensory response was indeed graded by visuo-tactile disparity (F = 6.02, p < 0.001, **Figure 2C**), being strongest when touch and vision were presented near each other and progressively decreasing with distance between them (D1 trough D5 respectively; 0.46μV ± 0.10μV, 0.34μV ± 0.11μV, 0.15μV ± 0.13μV, 0.06μV ± 0.08μV, −0.02μV ± 0.07μV). Most importantly, at near distances (D1 & D2, corresponding to 13.2 and 23.1cm) this metric had positive values significantly different from zero (all p < 0.006), indicating a true multisensory effect. This effect was present in controls (all p < 0.01) and in ASD individuals, but only for the nearest distance in the latter group (D1, p = 0.04; D2, p = 0.12). Thus, while the behavioral results failed to indicate differently sized PPSs in controls and ASD, the EEG suggests that these two groups do indeed differ in overall size (Mul et al., 2019; Noel et al., 2020).

The main effect differentiating controls (0.27μV ± 0.12μV) and ASD individuals (0.12μV ± 0.11μV) regardless of distance or social context approached but did not reach significance (p = 0.06), as did the contrast between social contexts regardless of experimental group and distance (p = 0.13). Critically, however, the group by social context did show a significant interaction (F = 3.74, p = 0.019). This latter effect was driven by the fact that social context altered the general degree of multisensory integration in neurotypical controls (non-social, 0.42μV ± 0.12μV; social, 0.14μV ± 0.16μV, p = 0.014), but not in individuals with ASD (non-social, 0.15μV ± 0.13μV; social, 0.14μV ± 0.13μV, p = 0.83). The 3-way interaction was not significant (p = 0.32).

### Neural Network Modeling: Differences in E/I balance in Autism may cause PPS inflexibility

Having established that both control and individuals with ASD showed a PPS effect behaviorally, as well as the presence of multisensory integration as indexed by physiology (i.e., superadditivity) that was greatest in the near space, we attempted to provide a neural modeling framework capable of accounting for the apparent inflexibility of PPS in ASD. For this purpose, we adapted an existing biologically-plausible neural network model of PPS (Magosso et al., 2010a, 2010b) by (i) reducing the number of neurons simulated, (ii) rendering it deterministic by eliminating noise sources, and (iii) introducing a mapping between neural activation and reaction times (see *Methods* and **Figure 3A** for details). Importantly, these three ingredients (and the targeted, hypothesis-driven fixing or not of parameters) allowed for the first time to use this neural network within the context of a model fitting procedure (Bogacz & Cohen, 2004, see **Supplementary Note** for detail).

First, we fitted the neural network to the psychophysical data from neurotypical individuals under the non-social condition. Here, *K*_1_, *K*_2_, *α*, and *Lim* (see **Eq. 3** and **Eq. 4**) were allowed to vary while the rest of parameters (**Table S1**) were fixed to the values in Magosso et al., 2010. These parameters were chosen given that they bear relation to features that are anatomical, likely immutable across the duration of the experiment but that could account for differences between individual participants. Most importantly for this first fit, results demonstrated that we were able to closely reproduce the pattern of reaction times exhibited by healthy controls during the nonsocial condition (RMSE = 0.41, **Figure 3B**). For the following steps, *K*_1_, *K*_2_, *α*, and *Lim* are set to this configuration, which serves as baseline.

Next, we attempted to account for the shrinking of PPS in control individuals during the social context (see findings here and in Teneggi et al., 2013). We entertained two possibilities. First, we fit the neural network with feedback synaptic weight (parameter 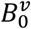, **Eq. 4**) as a free parameter, given that previous reports (Magosso et al., 2010a; Noel et al., 2020a) have postulated that changes in the strength of the synapses connecting unisensory and multisensory areas may account for the plastic resizing of PPS. However, the above possibility would require a very quick update in synaptic strengths, and thus we posited that instead the social context may reshape PPS by directly modulating the gain of the multisensory neuron (parameter *r^m^*, **Eq. 7**). This second potential source of modulation could originate from a number of long-range sources, for example, social cognition structures such as the amygdala or orbitofrontal cortex (see Clery et al., 2015, and Bufacchi & Iannetti, 2018 for a similar argument), and be implemented via a number of (quick) functional properties, such as altering the local chemical balance via neuromodulators. Model fits (**Figure 3B**) showed that the latter was the most likely possibility, with modification of *r^m^* (RMSE = 0.76) providing a better fit than alterations of 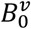 (RMSE = 2.03). Thus, the resizing of PPS during a social context is most parsimoniously explained by a modulation in the gain of the multisensory neuron.

As a last and most important step, we must explain why a social context – via modulation of *r^m^* – does not remap PPS in ASD. To do so, we considered that the E/I regime imposed by the “Mexican-hat” (within unisensory areas neurons are laterally connected by a “Mexican-Hat” pattern, with near excitation and far inhibition - see inset to the top of **Figure 3A** and **Eq. 2**) may be different in control neurotypical and ASD individuals. Indeed, a substantial literature suggests this possibility (Rubenstein & Merzenich, 2003; Lee et al., 2017), and simulations showed that increases in 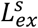 (i.e., more excitation) led to reductions in the size of PPS (**Figure S1**). That is, an increase in the relative strength of lateral excitations (vs. inhibition) of connections within sensory areas, as is hypothesized in ASD, would led to smaller PPS, which is what Mul et al., 2019, and Noel et al., 2020, have reported, and is numerically (but not statistically) consistent with the findings here. Thus, we let 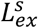 be a free parameter, and fit the ASD data during the non-social condition (starting from the baseline, control non-social model). This exercise improved the fit to data from the baseline neural network (RMSE = 0.54) and suggested a four-fold increase (from 0.15 at baseline to 0.63) in E/I ratio from neurotypical to individuals with ASD (**Figure 3C**). Most importantly, we then examined the potential impact *r^m^* would have under this new E/I regime. As alluded to above, we observed that under the default (control) condition, modulations of *r^m^* are reflected in the size of PPS (**Figure 3D**). However, under an elevated general state of lateral excitation, *r^m^* had little to no impact on the size of PPS (**Figure 3E**). In other words, the well-established E/I imbalance present in ASD (Rubenstein & Merzenich, 2003; Lee et al., 2017) can account for a weakening of the social signal related to the presence of another person, and thus renders the mapping of PPS inflexible in ASD.

## Discussion

We used a multisensory task where touch was applied on the body and visual stimuli were presented at different distances from neurotypical observers and individuals with ASD to map their PPS; this space being argued to implicitly index self-location and the division between self and other (Blanke, 2012; Serino, 2019; Noel et al., 2015). In line with results from Teneggi and colleagues (2013), PPS became smaller in neurotypical individuals within a social context. In previous work (Teneggi et al., 2013), this reduction in PPS size during a social condition has been interpreted as “giving space” to an unknown confederate (in Teneggi et al., 2013, this space then expands as to “include” the confederate after a positive social interaction). In the current study, we additionally describe for the first time the neural correlates of this social remapping of PPS. In line with observations made during intracranial recordings (Bernasconi et al., 2018), a neurophysiological “signature” of PPS could be seen as evidenced by changes in multisensory integration (i.e., difference between paired and summed responses) as a function of distance. Here we show that indeed this physiological marker of PPS is modulated by social context in neurotypical individuals. More importantly, both the behavioral and electrophysiological measures concurred in suggesting that PPS remapping and changes in multisensory integration due to social context did not occur in individuals with ASD.

The physiological recordings demonstrated superadditivity during multisensory presentations both in neurotypical participants and in individuals with ASD. This fact seemingly suggests that the basic processes of multisensory integration are intact in individuals with ASD, as recently suggested by behavioral reports focused on computational principles of behavior (Zaidel et al., 2015; Noel et al., 2020b). Instead, the results highlight a more specific anomaly; the modulation of multisensory integration, or lack thereof, in ASD via or during a social context. This finding adds to a series of studies emphasizing anomalies in the finer grain details of how individuals with ASD integrate information across sensory modalities, and adds to it through its emphasis on the spatial domain, as opposed to the much better studied temporal domain (Wallace et al., 2019; Stevenson et al., 2014; Baum et al., 2015; Noel et al., 2018c; Woynaroski et al, 2013). Indeed, alterations in fine grain multisensory temporal acuity in ASD (see Feldman et al., 2018, for a meta-analysis showcasing 53 studies in this domain), which have largely been studied for audio-visual pairings (vs. e.g., visuo-tactile), have been closely associated with language deficits (Bahrick & Todd, 2012; Woynaroski et al, 2013; Patten et al., 2014; Cascio et al., 2016). On the other hand, the focus of the current study was on visual-tactile pairings, where the range of biologically plausible temporal asynchronies is much more limited, given that touch necessarily occurs on the body. Thus, instead of the traditional focus on temporal alignment of multisensory signals, multisensory pairings involving the tactile modality are thought to subserve impact prediction (Clery et al., 2015b, 2018; Noel et al., 2018d), defensive (Graziano & Cooke, 2006), and approach-related (de Vignemont & Iannetti, 2015) behaviors. As such, multisensory pairings involving the tactile modality (e.g., visuo-tactile, audio-tactile) are likely to be more relevant for spatial coding, such as the construction and maintenance of PPS. In turn, PPS likely plays an important role in supporting non-verbal social behaviors (Noel et al., 2017), which represent a core feature in ASD that has been less studied relative to language deficits and their relation to (multi)sensory processing.

A critical contribution in the current report is the ability to perform formal model fitting of a biologically plausible neural network model of PPS. Namely, a number of prior reports have suggested potential changes in underlying neural circuitry to account for observed remapping of PPS. For instance, a popular suggestion is that remapping of PPS may occur due to changes in the strength of feedforward/feedback synapses via Hebbian Learning (e.g., Serino et al., 2015a; Noel et al., 2020a). And indeed, simulations suggest that in principle this could be the case. However, there are a number of other parameters within current network, in addition to the strength of synapses, that could resize PPS. One candidate is the gain of multisensory neurons. Thus, the framing of the neural network of PPS (Magosso et al., 2010a, b) under a model fitting procedure allows contrasting different hypotheses as to the mechanism that most likely supports the remapping of PPS. Here we find that in neurotypical individuals and during a social context manipulation it is not the strength of long-range synapses that best explains observed changes in the size of PPS (as it may in explaining the effect of tool use on PPS; Serino et al., 2015), but instead the gain of multisensory neurons (i.e., the steepness of their input-output relation, see **Figure 3A**) best accounts for empirical observations. In future work it will be interesting to examine whether the fitting of the neural network is sensitive to fine grain details of the re-sizing of PPS during different conditions yielding similar phenotypic results (e.g., enlargement of PPS during tool use or changes in perceived self-location; Serino et al., 2015 and Noel et al., 2015, respectively) and able to ascribe these effects that are similar at a surface level, to different underlying causes.

Collectively, our behavioral and electrophysiological results point to PPS being immune to social context in ASD. Our neural network model suggests that potential differences between the neurotypical control and ASD groups in regards to PPS resizing may be explained by well-known anomalies in E/I balance (Rubenstein & Merzenich, 2003; Lee et al., 2017), where the gain mechanism hypothesized to resize PPS is not effective under a regime of heightened excitation. These modeling results need to be confirmed experimentally. However, this framework nicely ties together a number of recent observations within the computational study of ASD. For instance, Rosenberg et al., 2015, were able to account for a number of perceptual deficits in ASD by postulating an anomaly in divisive normalization. This latter computation is ubiquitous in the central nervous system (Carandini & Heeger, 2011) and in essence amounts to contextualizing all output from a given neural area before impacting downstream targets (i.e., neural output from individual neurons are divided by the activity of a normalizing pool). In other words, there is a strong parallel between divisive normalization and changes in E/I balance provided by “Mexican-hat” lateral connections within a neural area. Perhaps even more interesting, Lieder and colleagues (2019) used an auditory frequency discrimination task to demonstrate that individuals with ASD showed a slow updating of Bayesian priors. Similarly, Lawson and colleagues (2017) suggested that adults with ASD overestimate the volatility of their sensory environment, and thus are slower in updating Bayesian priors when confronted with statistically unlikely events. In this line, Noel and colleagues (2018d) suggested that PPS is best conceived as a visuo-proprioceptive prior. Thus, the current findings agree with Bayesian accounts of ASD in suggesting an inflexible updating of priors (in this case, the prior being one of visuo-proprioceptive coupling). Further, by using a biologically-plausible neural network of PPS we are able to extend these observations and postulate a potential neural mechanism for this reduced flexibility. Thus, in the current work we bridge between Bayesian accounts of ASD suggesting reduced flexibility of priors (Lawson et al, 2017; Lieder et al., 2019), and circuit-level accounts suggesting an impairment in divisive normalization and/or excitation-inhibition imbalance (Rosenberg et al., 2015).

To conclude, here we attempt to link between social and sensory differences in ASD, and to add to the rapidly expanding field of computational psychiatry. We show via behavior and physiology that PPS updating is inflexible in ASD, which is reminiscent of statistical inference accounts suggesting a reduced flexibility of Bayesian priors (Lawson et al, 2017; Lieder et al., 2019). Additionally, we suggest a potential mechanism for this reduced flexibility – ineffective gain modulation within the E/I regime that has been described in ASD (Rubenstein & Merzenich, 2003; Lee et al., 2017). Broadly, given the role of PPS in bodily self-consciousness (Blanke, 2012), the current results may re-cast ASD as a disorder of the self – as it was originally described (Kanner, 1943) – and suggest how we may bridge across implementation and computational levels of description within the ASD pathology.

## References

Bahrick, L.E., Todd, J.T., 2012. Multisensory processing in autism spectrum disorders: intersensory processing disturbance as atypical development. In: Stein, B.E. (Ed.), The New Handbook of Multisensory Processes. MIT Press, Cambridge, MA, pp. 657–674.

Baum, S. H., Stevenson, R. A. & Wallace, M. T. (2015). Behavioral, perceptual, and neural alterations in sensory and multisensory function in autism spectrum disorder. Prog. Neurobiol 134, 140–160.

Bausenhart, K. M., Di Luca, M., & Ulrich, R. (2018). Assessing duration discrimination: Psychophysical methods and psychometric function analysis. In Timing and Time Perception: Procedures, Measures, & Applications (pp. 52–78): Brill.

Bernasconi, F., Noel, J.P., Park, H.D., Faivre, N., Seeck, M., Spinelli, L., Schaller, K., Blanke, O., Serino, A. (2018). Spatio-temporal processing of multisensory peripersonal space in human parietal and temporal cortex: an intracranial EEG study. Cerebral Cortex. https://doi.org/10.1093/cercor/bhy156

Blanke, O. (2012). Multisensory brain mechanisms of bodily self-consciousness. Nat. Rev. Neurosci. 13, 556–571. doi: 10.1038/nrn3292

Bogacz, R., Cohen, J.D. (2004). Parameterization of connectionist models, Behavior Research Methods, Instruments, & Computers 36(4) 732–741.

Bufacchi, R.J.; Iannetti, G.D. An action field theory of peripersonal space. Trends Cogn Sci. 2018, 22, 1076–1090.

Cappe, C., Thut, G., Romei, V., & Murray, M. M. (2010). Auditory-visual multisensory interactions in humans: Timing, topography, directionality, and sources. Journal of Neuroscience, 30, 12572–12580

Carandini M, Heeger DJ. (2011). Normalization as a canonical neural computation. Nat. Rev. Neurosci. 13(1):51–62

Cascio, C.J., Gu, C., Schauder, K.B., Key, A.P., & Yoder, P. (2015). Somatosensory event-related potentials and association with tactile behavioral responsiveness patterns in children with ASD. Brain Topography, 28(6), 895–903.

Cascio, C.J., Woynaroski, T., Baranek, G.T., Wallace, M.T., (2016). Toward an interdisciplinary approach to understanding sensory function in autism spectrum disorder. Autism Res. 9, 920–925. https://doi.org/10.1002/aur.1612.

Chen, Q., Deister, C.A., Gao, X., Guo, B., Lynn-Jones, T., Chen, N. et al., (2020). Dysfunction of cortical GABAergic neurons leads to sensory hyper-reactivity in a Shank3 mouse model of ASD. Nature Neuroscience; DOI: 10.1038/s41593-020-0598-6

Cléry, J. & Hamed, S. B. (2018). Frontier of Self and Impact Prediction. Frontiers in Psychology 9

Cléry, J., Guipponi, O., Odouard, S., Wardak, C., and Ben Hamed, S. (2015b). Impact prediction by looming visual stimuli enhances tactile detection. J. Neurosci. 35, 4179–4189. doi: 10.1523/JNEUROSCI.3031-14.2015

Cléry, J., Guipponi, O., Wardak, C., and Ben Hamed, S. (2015a). Neuronal bases of peripersonal and extrapersonal spaces, their plasticity and their dynamics: knowns and unknowns. Neuropsychologia 70, 313–326. doi: 10.1016/j.neuropsychologia.2014.10.022

Constantino, J. N., Davis, S. A., Todd, R. D., Schindler, M. K., Gross, M. M., Brophy, S. L., … Reich, W. (2003). Validation of a brief quantitative measure of autistic traits: Comparison of the social responsiveness scale with the autism diagnostic interview revised. Journal of Autism and Developmental Disorders, 33, 427–433. https://doi.org/10.1023/A:1025014929212.

de Vignemont, F., and Iannetti, G. D. (2015). How many peripersonal spaces? Neuropsychologia 70, 327–334. doi: 10.1016/j.neuropsychologia.2014.11.018

Delorme, A., & Makeig, S. (2004). EEGLAB: An open source toolbox for analysis of single-trial EEG dynamics including independent component analysis. Journal of Neuroscience Methods, 134, 9–21

Drugowitsch, J., DeAngelis, G.C., Klier, E.M., Angelaki, D.E. and Pouget, A. (2014). Optimal multisensory decision-making in a reaction-time task. Elife, 3.

Feldman, J. I., Dunham, K., Cassidy, M., Wallace, M. T., Liu, Y., & Woynaroski, T. G. (2018). Audiovisual multisensory integration in individuals with autism spectrum disorder: A systematic review and meta-analysis. Neuroscience and Biobehavioral Reviews, 95, 220–234

Gogolla, N., Takesian, A. E., Feng, G., Fagiolini, M. & Hensch, T. K. (2014). Sensory integration in mouse insular cortex reflects GABA circuit maturation. Neuron 83, 894–905 (2014).

Goncalves, J. T., Anstey, J. E., Golshani, P. & Portera-Cailliau, C. (2013). Circuit level defects in the developing neocortex of Fragile X mice. Nat. Neurosci. 16, 903–909

Graziano, M., & Cooke, D. (2006). Parieto-frontal interactions, personal space, and defensive behavior. Neuropsychologia, 44(13), 2621–2635.

Guthrie, D., & Buchwald, J. S. (1991). Significance testing of difference potentials. Psychophysiology, 28, 240–244.

Hahnloser, R., Sarpeshkar, R., Mahowald, M. A., Douglas, R. J., Seung, H. S. (2000). Digital selection and analogue amplification coexist in a cortex-inspired silicon circuit. Nature, 405 (6789): 947–951

Happé F., Frith U. (2006). The weak coherence account: Detail-focused cognitive style in autism spectrum disorders. Journal of Autism and Developmental Disorders.;36:5–25

Hobeika L, Taffou M, Carpentier T, Warusfel O, Viaud-Delmon I. (2019). Capturing the dynamics of peripersonal space by integrating expectancy effects and sound propagation properties. J Neurosci Methods. 2019 Dec 2;332:108534

Huys, Q. J. M., Maia, T. V., & Frank, M. J. (2016). Computational psychiatry as a bridge from neuroscience to clinical applications. Nature Neuroscience, 19(3), 404–413.

Ishida, H., Nakajima, K., Inase, M., Murata, A., (2010). Shared mapping of own and others’ bodies in visuotactile bimodal area of monkey parietal cortex. J Cogn Neurosci 22, 83–96. doi:10.1162/jocn.2009.21185

Joseph RM, Keehn B, Connolly C, Wolfe JM, Horowitz TS. 2009. Why is visual search superior in autism spectrum disorder? Developmental Science 12: 1083–96

Kandula, M., Van der Stoep, N., Hofman, D., and Dijkerman, H. C. (2017). On the contribution of overt tactile expectations to visuo-tactile interactions within the peripersonal space. Exp. Brain Res. 235, 2511–2522. doi:10.1007/s00221-017-4965-9

Kanner L. (1943). Autistic disturbances of affective contact. Nervous Child;2:217–250.

Karvelis, P., Seitz, A., Lawrie, S., Series, P. (2018). Autistic traits, but not schizotypy, predict overweighting of sensory information in Bayesian visual integration, eLife, 7:e34115.

Làdavas, E., Zeloni, G. & Farnè, A. (1998). Visual peripersonal space centred on the face in humans. Brain 121, 2317–2326.

Lawson RP, Mathys C, Rees G (2017) Adults with autism overestimate the volatility of the sensory environment. Nat Neurosci 20:1293–1299

Lee E, Lee J, Kim E. Excitation/inhibition imbalance in animal models of autism spectrum disorders. Biol Psychiatry. 2017;81(10):838–47.

Lehmann, D., & Skrandies, W. (1980). Reference-free identification of components of checkerboard evoked multichannel potential fields. Electroencephalography and Clinical Neurophysiology, 48, 609–621

Lieder, I., Adam, V., Frenkel, O., Jaffe-Dax, S., Sahani, M., & Ahissar, M. (2019). Perceptual bias reveals slowupdating in autism and fast-forgetting in dyslexia. Nature Neuroscience, 22(2), 256

Lord C, Rutter M, DiLavore PC, Risi S, Gotham K, Bishop S. Autism Diagnostic Observation Schedule, Second Edition (ADOS-2) Manual (Part I): Modules 1-4. Torrance, CA: Western Psychological Services; 2012.

Magosso E, Ursino M, di Pellegrino G, Ladavas E, Serino A. (2010a). Neural bases of peri-hand space plasticity through tool-use: insights from a combined computational-experimental approach. Neuropsychologia 48: 812–830. doi:10.1016/j.neuropsychologia.2009.09.037.

Magosso E, Zavaglia M, Serino A, di Pellegrino G, Ursino M. (2010b) Visuotactile representation of peripersonal space: a neural network study. Neural Comput 22: 190–243. doi:10.1162/neco.2009.01-08-694.

Masson, C. J., van der Westhuizen, D., Noel, J., Prevost, A. M., van Honk, J., Fotopoulou, A., … Serino, A. (2020). Testosterone administration in women increases the size of their peripersonal space. PsyArXiv https://doi.org/10.31234/osf.io/8cmtp

Miller J. 1982. Divided attention: evidence for coactivation with redundant signals. Cognitive Psychology 14:247–279. doi: 10.1016/0010-0285(82)90010-X.

Mul, C. L., Cardini, F., Stagg, S. D., Sadeghi Esfahlani, S., Kiourtsoglou, D., Cardellicchio, P., et al. (2019). Altered bodily self-consciousness and peripersonal space in autism. Autism, 1362361319838950.

Murray, M. M., Brunet, D., & Michel, C. M. (2008). Topographic ERP analyses: A step-by-step tutorial review. Brain Topography, 20, 249–264.

Noel, J-P. Bertoni, T., Pellenin, E., Herbelin, B., Magosso, E., Blanke, O., Wallace, M., Serino, A. (2020a). Rapid recalibration of peri-personal space; behavioral, electrophysiological, and computational evidence. Cerebral Cortex

Noel, J-P., Chatelle, C., Perdikis, S., Jöhr, J., Lopes Da Silva, M., Ryvlin, P., De Lucia, M., del R. Millán, J., Diserens, K., Serino, A., (2019). Peri-personal space encoding in patients with disorders of consciousness and cognitive-motor dissociation. NeuroImage: Clinical

Noel, J-P., Lakshminarasimhan, K., Park, H., Angelaki, D. (2020b). Increased variability but intact integration during visual navigation in Autism Spectrum Disorder. Proceedings of the National Academy of Science

Noel, J. P., Samad, M., Doxon, A., Clark, J., Keller, S., & Di Luca, M. (2018e). Peri-personal space as a prior in coupling visual and proprioceptive signals. Scientific Reports, 8, 15819. doi:10.1038/s41598-018-33961-3

Noel, J. P., Stevenson, R. A., & Wallace, M. T. (2018). Atypical audiovisual temporal function in autism and schizophrenia: Similar phenotype, different cause. European Journal of Neuroscience, 47, 1230–1241.

Noel, J.-P., Park, H.-D., Pasqualini, I., Lissek, H., Wallace, M., Blanke, O., & Serino, A. (2018). Audio-visual sensory deprivation degrades visuo-tactile peri-personal space. Consciousness and Cognition, 61, 61–75.

Noel, J.P., Blanke, O., Magosso, E. Serino, A. (2018b). Neural Adaptation Accounts for the Resizing of Peri-Personal Space Representation: Evidence from a Psychophysical-Computational Approach. Journal of Neurophysiology. https://doi.org/10.1152/jn.00652.2017

Noel, J.P., Cascio, C., Wallace, M., Park, S. (2017). The spatial self in Schizophrenia and Autism Spectrum Disorder. Schizophrenia Research, 170, 8–12

Noel, J.P., Failla, M.D., Quide-Zlibut, J.M., Williams, Z.J., Gerdes, M., Trace, J.M., Zoltowski, A. R., Foss-Feig, J.H., Nichols, H. S., Armstrong, K., Heckers, S.H., Blake, R., Wallace, M.T., Park, S. Cascio, C. (2020). Visual-tactile spatial multisensory interaction in adults with autism and schizophrenia. Front. Psychiatry, doi: 10.3389/fpsyt.2020.578401

Noel, J.P., Pfeiffer, C., Blanke, O., Serino, A. (2015). Full body peripersonal space as the sphere of the bodily self? Cognition, 144 (3), 49–57.

Noel, J.P., Serino, A., Wallace, M. (2018a). Increased neural strenght and reliability at the boundary of peripersonal space. Journal of Cognitive Neuroscience, doi: 10.1162/jocn_a_01334

Noel, J. P., De Niear, M., Lazzara, N. S., & Wallace, M. T. (2017). Uncoupling between multisensory temporal function and nonverbal turn-taking in autism spectrum disorder. IEEE Transactions on Cognitive and Developmental Systems. https://doi.org/10.1109/TCDS.2017.2778141

Orefice, L. L. et al. (2016). Peripheral mechanosensory neuron dysfunction underlies tactile and behavioral deficits in mouse models of ASDs. Cell 166, 299–313

Palmer CJ, Lawson RP, Hohwy J. (2017). Bayesian approaches to autism: Towards volatility, action, and behavior. Psychol Bull.;143: 521–542.

Patten, E., Watson, L.R., Baranek, G.T., 2014. Temporal synchrony detection and associations with language in young children with ASD. Autism Res. Treat. 2014, 1–8. https://doi.org/10.1155/2014/678346

Pellencin, E., Paladino, M. P., Herbelin, B., & Serino, A. (2018). Social perception of others shapes one’s own multisensory peripersonal space. Cortex, 104, 163–179.

Pellicano, E. & Burr, D. (2012). When the world becomes ‘too real’: a Bayesian explanation of autistic perception. Trends Cogn. Sci. 16, 504–510

Perrin, F., Pernier, J., Bertnard, O., Giard, M. H., & Echallier, J. F. (1987). Mapping of scalp potentials by surface spline interpolation. Electroencephalography and Clinical Neurophysiology, 66, 75–81

Pfeiffer, C., Noel, J.P., Blanke, O., Serino, A. (2018). Vestibular modulation of peri-personal space boundaries. European Journal of Neuroscience, 47, 800–811, doi:10.1111/ejn.13872

Pitkow X, Angelaki DE: Inference in the brain: statistics flowing in redundant population codes. Neuron 2017, 94:943–953.

Rizzolatti, G., Scandolara, C., Matelli, M., Gentilucci, M., (1981). Afferent properties of periarcuate neurons in macaque monkeys. II. Visual responses. Behav. Brain Res. 2, 147–163.

Rizzolatti, G., Matelli, M., Pavesi, G., (1983). Deficits in attention and movement following the removal of postarcuate (area 6) and prearcuate (area 8) cortex in macaque monkeys. Brain 106 (3), 655–673.

Robertson, C. E. & Baron-Cohen, S. (2018). Sensory perception in autism. Nature Reviews Neuroscience, 18(11), 671.

Rosenberg A, Patterson JS, and Angelaki DE (2015) A computational perspective on autism. Proceedings of the National Academy of Sciences, 112(30): 9158–9165

Rubenstein JL, Merzenich MM. Model of autism: increased ratio of excitation/inhibition in key neural systems. Genes Brain Behav. 2003;2(5):255–67.

Salomon, R., Noel, J.P., Lukowska, M., Faivre, N., Metzinger, T. Serino A., Blanke, O. (2017). Unconscious visuo-tactile integration shapes peripersonal space representation and modulates bodily self-consciousness. Cognition, 166, 174–183

Serino A (2019) Peripersonal space (PPS) as a multisensory interface between the individual and the environment, defning the space of the self. Neurosci Biobehav Rev 99:138–159

Serino A, Canzoneri E, Marzolla M, di Pellegrino G, Magosso E. (2015a) Extending peripersonal space representation without tool-use: evidence from a combined behavioral-computational approach. Front Behav Neurosci 9: 4, doi:10.3389/fnbeh.2015.00004.

Serino, A., Noel, J.P., Galli, G., Canzoneri, E. Marmaroli, P., Lissek, H., Blanke, O. (2015b). Body part-centered and full body-centered peripersonal space representations. Scientific reports, vol. 5, p. 18603.

Shelton, P. A., Bowers, D., & Heilman, K. M. (1990). Peripersonal and vertical neglect. Brain, 113, 191–205. https://doi.org/10.1093/brain/113.1.191.

Simon, D. M., Noel, J. P., & Wallace, M. T. (2017). Event related potentials index rapid recalibration to audiovisual temporal asynchrony. Frontiers in Integrative Neuroscience, 11, 8

Sperdin HF, Cappe C, Murray MM (2010) The behavioral relevance of multisensory neural response interactions. Front Neurosci 4:9.

Stein, B.E., Stanford, T.R. (2008). Multisensory integration: current issues from the perspective of the single neuron. Nat. Rev. Neurosci. 9, 255–266

Stevenson, R. A., Siemann, J. K., Schneider, B. C., Eberly, H. E., Woynaroski, T. G., Camarata, S. M., et al. (2014). Multisensory temporal integration in autism spectrum disorders. The Journal of Neuroscience, 34(3), 691–697

Teneggi, C., Canzoneri, E., di Pellegrino, G., & Serino, A. (2013). Social modulation of peripersonal space boundaries. Current biology, 23(5), 406–411.

Virtanen, P., Gommers, R., Oliphant, T. E., Haberland, M., Reddy, T., Cournapeau, D., Burovski, E., Peterson, P., Weckesser, W., Bright, J. et al., (2020). Scipy 1.0: fundamental algorithms for scientific computing in python, Nature methods 17 (3) 261–272.

Wallace, M., Woynaroski, T, Stevenson, RA (2019). Multisensory integration as a window into orderly and disrupted cognition and communication. Annual review of Psychology, 71

Wormington, M. Panaccione, C., Matney, K.M., Bowen, D.K. (1999). Character-ization of structures from x-ray scattering data using genetic algorithms, Philosophical Transactions of the Royal Society of London. Series A: Mathematical, Physical and Engineering Sciences 357 (1761) 2827–2848.

Woynaroski, T.G., Kwakye, L.D., Foss-Feig, J.H., Stevenson, R.A., Stone, W.L., Wallace, M.T. (2013). Multisensory speech perception in children with autism spectrum disorders. J. Autism Dev. Disord. 43, 2891–2902. https://doi.org/10.1007/s10803-013-1836-5.

Xu, G., Strathearn, L., Liu, B., O’Brien, M., Kopelman, T. G., Zhu, J., … Bao, W. (2018). Prevalence and treatment patterns of autism spectrum disorder in the United States, 2016. JAMA Pediatrics, 173(2), 153–159. https://doi.org/10.1001/jamapediatrics.2018.4208

Zaidel A, Goin-Kochel RP, Angelaki DE (2015) Self-motion perception in autism is compromised by visual noise but integrated optimally across multiple senses. Proc Natl Acad Sci USA 112(20):6461–6466

